# Observations of Elemental Composition of Enceladus Consistent with Generalized Models of Theoretical Ecosystems

**DOI:** 10.1101/2023.10.29.564608

**Authors:** Daniel Muratore, Sara I. Walker, Heather Graham, Christopher H. House, Christopher P. Kempes

## Abstract

Recent analysis of data from the Cassini Cosmic Dust Analyzer confirm geochemical modeling work that shows that the ocean of Enceladus contains considerable quantities of inorganic phosphorus as well as ammonium [55]. Technological advancement in flight instrumentation will continue to yield increasingly detailed data about the relative elemental and molecular composition of life detection candidates. Apart from speculating about threshold concentrations of bioactive compounds to support ecosystems, metabolic and ecological theory can provide a powerful interpretative lens to assess whether extraterrestrial environments are compatible with living ecosystems. Using multiple levels of ecological analysis, spanning from assuming strictly Earth-like organismal physiology to more agnostic understandings of putative biochemistries, we compare the proposed N:P stoichiometries of the Enceledus ocean to possible ecologies. We use chemostat models to predict potentially supportable biomass concentrations under different energy and matter flux regimes, macromolecular allometric theory to compare hypothetical biomass N:P ratios to possible environmental N:P supply ratios, and finally take a data-driven biogeochemical approach to predict possible biomass N:P ratios from the observed dissolved ratios. All three of our modeling approaches suggest marginal compatibility of an ecosystem with the ranges of dissolved N and P concentrations in the Enceledean ocean. Based on our analysis, we suggest two main priorities for further research into terrestrial analogs to improve our ability to interpret geochemical ratios as a life detection instrument.

## Introduction

One the most consistent patterns associated with life on Earth is the Redfield ratio of elements [19, 28, 58]. Originally discovered in particulate organic matter in seawater [58], this ratio describes the abundances of three of life’s essential elements, relating carbon to nitrogen to phosphorus in a ratio of 106:16:1. This ratio is found across the global ocean, in environments ranging from low to high biomass, and across the tree of life [2, 66, 67]. Because of this seeming ubiquity, the Redfield ratio has been considered a target signature for astrobiological life detection, especially on ocean worlds such as Europa and Enceladus [2, 11, 15, 39, 46, 73, 78]. Furthermore, these elemental ratios are consistent across a diverse range of ecosystems [66, 67], allowing for their use in the case where precise estimates of elemental or molecular abundances can be technologically challenging. Analysis of elemental ratios on life detection candidates can also facilitate mechanistically principled reasoning about the relationships between abiotic resource and hypothetical biotic elemental compositions.

Recent data from the Cassini Cosmic Dust Analyzer (CDA), reporting mass spectra of Type 3 (Na-salt-containing) water ice grains from the plume of Enceladus have been used to assert high concentrations of inorganic phosphate in the Enceladus ocean [55]. Observations follow geochemical modeling simulations that also suggest significant concentration These observations follow from geochemical simulations that also suggest significant concentrations of phosphate [23]. These reports of phosphorus follow on the tails of previous work identifying numerous elemental constituents of terrestrial life (C, N, H, O) from the Enceladus plume [53, 72]. Futher, analysis of higher molecular-weight mass spectra have suggested the presence of molecules common in living organisms, such as amino acid precursors, carboxylates, ammonium, and hydrocarbons from the plume [30, 54]. The ice grains analyzed by the CDA are typically sub-micron in size, and due to the high salt content (emblematized by a large Na+ mass line) in Type 3 particles, we assume that the concentrations predicted from these particles more accurately represent the bulk solution ejected into the plume as opposed to possible cellular debris - indicating these molecules are likely dissolved in Enceladus seawater and putatively available for biological uptake and utilization. The observation of phosphorus as a constituent of Enceladus seawater further contributes to the notion that the chemical components necessary for Earth-life are available and may be able to support biochemistry. While a metabolic network expansion study was unable to find an individual, completely sequenced terrestrial microbial genome capable of operating a full metabolism with the suite of compounds already confirmed to be detected in the Enceladus plume [60], these observations provide a basis for more seriously considering the possibilities of terrestrially-analogous biochemistry occurring in the Enceladus ocean.

In particular, due to the presence of both diatomic hydrogen and carbon dioxide in plume data in disequilibrium with the gas methane [5, 26, 42, 72], the hypothesis that Encladus may support methanogenesis (the biological production of methane as a metabolism) has emerged [1]. Methanogenesis is a metabolism performed by Archaea across a wide variety of Earth environments, including putative analogs to the Enceladus ocean [12, 56]. Using cultured methanogenic Archaea, laboratory studies have confirmed methanogenic growth under abiotic conditions that mirror those of Enceladus [68].

Biogeochemical modeling studies based on terrestrial methanogens have suggested compatibility of methanogenic Archaea with environments in the Enceladus ocean. Biomass estimates based on thermodynamic calculations of energy available for cellular growth have ranged from 10^5^ *−* 10^11^ cells per mL of Enceladus seawater or hydrothermal vent fluid (see Table 1). Using niche-based modeling, Tenelanda-Osorio et al. suggest potential biomass concentrations of 10^10^*−* 10^11^ cells/mL [69]. Using a metabolic model based on apparent amino acid concentrations, Steel et al. suggest cells growing in hydrothermal vent fluid at concentrations of 10^9^ cells/mL may dilute to 100 *−* 10000 cells/mL in the overlying seawater [64]. Affholder et al. take a Bayesian approach to compare the likelihood of an Enceladus with biological methanogenesis versus one without to observations of H_2_ : CO_2_ from Cassini and determine that methanogenesis, or environmental conditions that could facilitate methanogenesis, is a likely scenario. [1].

**Table 1.** Summary of putative biomass estimates using different energetics-based modeling techniques for Enceladus.

| Approach | Minimum Estimate | Maximum Estimate | Reference |
| --- | --- | --- | --- |
| Niche-Modeling | $10^{10}$ cells/mL | $10^{11}$ cells/mL | [69] |
| Metabolic Model | 100 cells/mL (ocean) | $10^9$ cells/mL (vent) | [64] |
| Geothermal Fluxes, Plume Optics | $10^5$ cells/mL | $10^8$ cells/mL | [52] |

Previous work emphasizing thermodynamics to model methanogenic metabolism on Enceladus have not taken into account resource uptake to synthesize biomass, and therefore cannot be further clarified with new and incoming data about dissolved nitrogen and phosphorus stocks. Instead, the dissolved concentrations of these elements have been compared to their concentrations on Earth environments [23, 42, 55], and because the concentrations are estimated to be much higher on Enceladus than Earth, have been considered to pass a threshold requirement for supporting life [55]. A biomass-focused ecosystem stoichiometric approach can provide further refinement to these arguments, as apparent energy utilization can be tied to expectations of elemental uptake and ecosystem macromolecular composition and ratios. Biomass composition can then be systematically related to observed dissolved inorganic ratios of putative nutrients as an additional test of the hypothesis that apparently energy utilization is tied to a system that assimilates resources into biochemical machinery.

We propose combining biogeochemical modeling with recent flight mass spectral data to include biomass composition in addition to metabolic energy flux in our suite of life detection approaches [1, 69]. Namely, we present: 1) chemostat models incorporating nutrients, 2) macromolecular allometric scaling for ecosystems with size distributions corresponding to putative terrestrial analogs, 3) generalized scaling of elemental ratios as a function of cellular physiology and ecosystem size-structure. Through analysis of our chemostat model, we can arrive at equilibrium biomass estimates and cellular turnover times that are comparable to previous energetics-based biogeochemical modeling efforts. Using macromolecular scaling theory, we can then pose a thought experiment estimating the elemental composition of an ecosystem comprised entirely of methanogenic Archaea to compare with observed inorganic N:P ratios, and generalize that thought experiment to consider hypothetical biochemistries obeying fundamentally different elemental scaling behavior to microbial life on Earth. This more agnostic approach also allows us to situate observed Enceladus ammonium:phosphate ratios in the context of Earth seawater chemistry, generating some hypotheses about the putative biological relevance of phosphorus to hypothetical Enceladus ecosystems. Applying these approaches to Enceladus as characterized by the Cassini mission yields three independent, yet mutually compatible, interpretations of the dissolved nitrogen and phosphorus ratios of the Enceladus ocean, each of which illustrates marginal compatibility with - and certainly does not rule out - a living ecosystem.

### Interpreting Environmental Ratios from Ecological Models

In order to interpret the observed elemental abundances of Enceladus and their possible consistency with a living world we need a way to solve for the steady-state of a coupled ecosystem and its environment. Understanding the biogeochemistry on Earth has successfully relied on simple chemostat - flow through system - models of coupled cellular physiology, abundance, and environmental nutrient concentrations. These models have recently been generalized for astrobiological contexts [28], and are an appropriate tool for basic understanding of the potential Enceladus geobiosphere.

We implemented a single-nutrient chemostat model similar to the model presented in [28] (see Methods). Briefly, our model assumes nitrogen to be replete (due to the high reported concentrations [42, 72]). In our chemostat setup we treat phosphorus as the limiting nutrient, adjust maximum growth rates according to a range possible energy fluxes, consider an average cell size (which dictates its physiological parameters), and use flow rates estimated from Enceladus (see Methods, Extended Data Figure 1). This model is a dual energy and resource limitation framework and allows us to estimate the expected biomass concentration consistent with the nutrient concentrations, physical parameters, and the possible energy regime of Enceladus.

Our model predicts ranges of steady-state biomass concentrations (Figure 1) given Enceladus conditions that match previous, independent estimates [52, 64, 69] and would be comparable to eutrophic environments on Earth [75] (see Extended Data Table 1). At the upper end of estimated P concentrations, we find ranges of estimates for higher growth efficiencies that match the estimates offered by [69] and [64] for life growing in hydrothermal fluid. Meanwhile, our low energy estimates match the biomass estimate of 10^5^ cells/mL from [52]. For the lowest phosphorus concentrations reported in [55], our estimated steady state cell densities range from approximately 10^2^ *−* 10^6^, bracketing the range of 10^5^ cells/mL estimated in [52] and 80-4250 cells/mL suggested in [64]. The Steel et al. model assumes that cells were only produced in the hydrothermal fluid and then diffusely mixed into the entire ocean volume [64]. Our biomass estimates are lower in the low phosphorus regime than the energetics-based models biomass estimates indicating that the phosphorus concentrations reported in [55] could in fact be limiting for a hypothetical Enceladus ecosystem.

**Figure 1.**
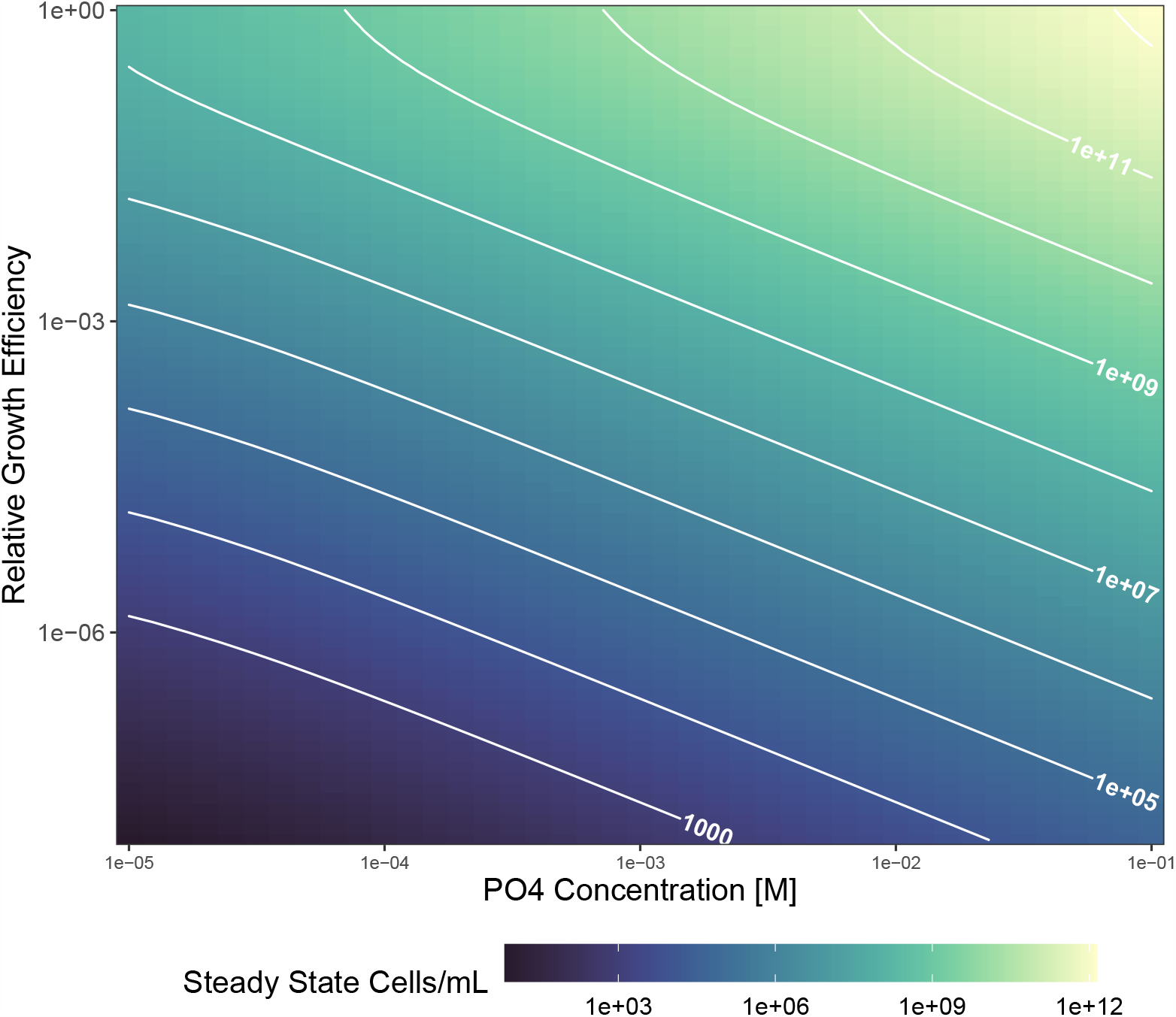
Chemostat model biomass calculations for Enceladus across possible parameter ranges. X-axis denotes steady state dissolved inorganic phosphate concentration, y-axis denotes relative growth efficiency of cells (with 1 being cell growth at rates comparable to a Terran analog). Note log scale for both axes. Color indicates steady state biomass in units of cells/mL. Contours indicate isolines of steady state biomass.

Simultaneously incorporating energy limitation and resource limitation in a chemostat model recapitulates previous estimates of possible sustainable biomass on Enceladus, but further refines those estimates through phosphorus limitation at lower phosphate concentrations. However, the basic chemostat model is limited in two ways: first, it does not account for the diversity of cellular physiology in an ecosystem. Second, it is strongly conditioned on the physiology of microorganisms on Earth. A more comprehensive assessment of a putative Encaledean biosphere should rely on complete ecosystem models and relax assumptions about the physiology of life, both of which we investigate in the following sections. We can more closely examine the details of cellular physiology and elemental composition through modeling techniques that leverage cellular macromolecular scaling laws, both by considering the more specific case of Earth methanogens and the more general case of physiologies not standard on Earth.

### Predicting Stoichiometries for a Hypothetical Methanogenic Ecosystem

Using size as a master trait to infer a variety of physiological features and ecological interactions [3] has been a successful approach in biogeochemical models [18, 74] and reconciling plankton diversity across the ocean [6, 7]. The most general version of this framework to tally biomass elemental totals is to combine the distribution of organism sizes with allometric scaling relationships for macromolecular components. While both cell size distributions and macromolecular allometries are general across life on Earth [9, 10, 29], they can be easily generalized to consider alternative physiologies and ecosystem structures [28]. We deploy a size-based framework to first generate elemental ratio expectations for Enceladus if the ecosystem were comprised entirely of methanogenic Archaea as known on Earth, we then generalize the perspective to life that radically departs from Earth-like parameters.

In order to generate ecosystem-level elemental abundances and ratios we first need to estimate a size distribution of cells. Nucleic acids are a class of macromolecule with established allometric scalings against cell size [28]. In particular, genome length has a well-described power-law scaling with cell volume [29] (see Extended Data Table 2). To simulate a hypothetical ecosystem comprised of methanogens, we used a compendium of genome sequences for known terrestrial methanogens [56]. We estimated a distribution of genome lengths for methanogenic Archaea based on closed genomes from 74 taxa spanning a broad range of temperature niches [61](see Methods). Water temperatures in the Enceladus ocean are estimated to be low in comparison to Earth surface temperatures (0^*°*^ C), so before proceeding with allometric conversions we confirmed that psychrotolerant Archaea did not systematically deviate from the genome length distribution of other methanogens (Figure 2A). Because they did not, we leveraged all available genomes from Prondzinksy et al. to estimate the genome length distribution. By inverting the genome size to cell size allometry, we were able to convert the genome length distribution into an approximate cell volume distribution (Figure 2B). We then fit a gamma distribution to the estimated cell volumes to impute a potential cell size distribution (see Methods). After calculating macromolecular scalings described below across the range of volumes (Figure 2C), we integrate over this size distribution to estimate ecosystem-level elemental and macromolecular composition (Figure 2D).

**Figure 2.**
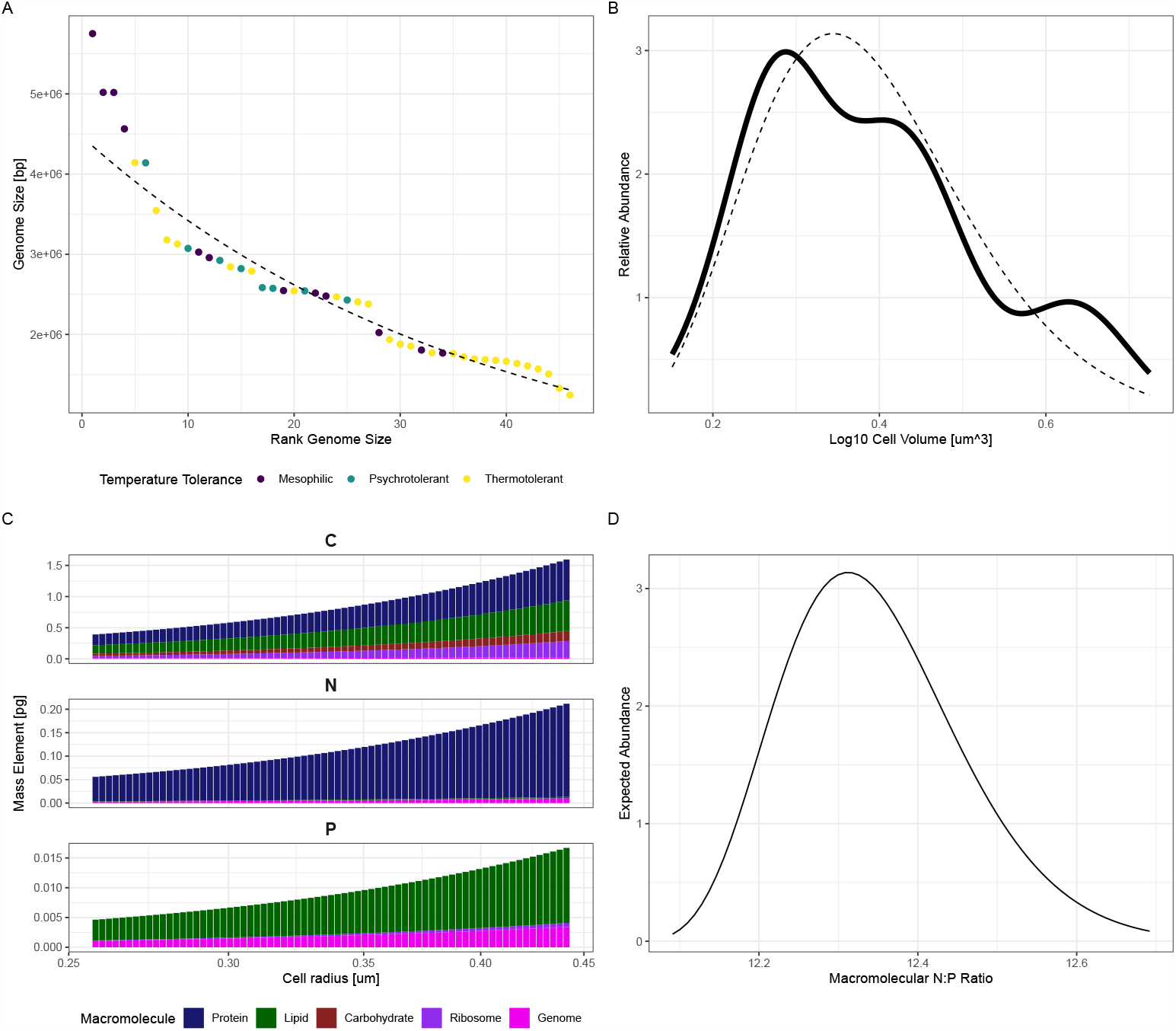
Elemental compositional analysis of sequenced Earth methanogens. A) Distribution of genome sizes of complete genome sequences for known methanogenic Archaea, data from [56]. Dashed line indicates exponential fit of rank-size distribution(Genome length (Mbp)= 4.5 * *e*^−0.0267\**r*^, 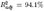), where r is the rank of the genome length. B) Distribution of inferred cell volumes (note log scale) based on lengths of complete genome sequences. Dashed line indicates gamma distribution fit to account for low sample size (∼ Γ(1606 ± 253, 252 ± 40)). C) Scaling of cellular carbon, nitrogen, phosphorus as a function of cell radius based on macromolecular allometries. Y-axis denotes the mass of each element in picograms for each cell radius (x-axis), the bar color denotes which macromolecule contributes to that element. D) Emergent distribution of cellular N:P ratios as a function of cell size (C) and relative abundance according to the estimated cell size distribution (B).

For this study, we compiled allometric scaling models that generalize well across bacteria and unicellular eukaryotic organisms measuring the pools of proteins, lipids, carbohydrates, ribosomes, and genomes as a function of cell size [17, 29]. While Archaea do have slightly different macromolecular architectures from Bacteria, such as tetraether membrane lipids [22], cellular stoichiometric data from extreme environments dominated by these organisms tend not to deviate far from Redfield elemental ratios [47], suggesting these macromolecular differences do not make a large difference in cellular elemental contents. These macromolecules are representative of cellular functions including information storage and processing, energy storage and utilization, biomass synthesis, and production of enzymes for biochemical potential, and these pools together account for most of the dry weight of terrestrial cells [38]. We then used average elemental compositions of each of these macromolecules (see Methods) to estimate cellular C, N, and P totals as a function of cell volume (Figure 2C). We analyzed the emergent elemental scalings as a result of the summed macromolecular scaling calculations. Our model predicts a slow scaling of N:P with respect to cell volume (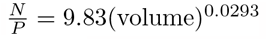), where volume is in cubic microns. This small exponent means that according to these models, N:P doesn’t change very much across small changes in cell size. Consequently, bulk ecosystem stoichiometries should be robust to variability in the cell size distribution for communities with a narrow range of cell sizes, such as the one we consider in this thought experiment.

We estimated the bulk ecosystem stoichiometry by integrating the product of each elemental scaling over the empirically-fitted cell volume distribution (see Methods), then taking their ratios. Under these conditions, the bulk community C:N:P composition is approximately 91:12:1 (see Figure 2D). This ratio is close to the canonical Redfield ratio, and is compatible with ratios measured from diverse ecosystems on Earth incorporating many metabolisms [24, 65–67]. Notably, we predict low N:P and C:P ratios relative to the Redfield ratio [19], possibly in part due to the small cell sizes of methanogenic organisms(e.g., [25]), or possibly due to not modeling photosynthetic pigments, macromolecules rich in C and N but low in P that underlie many food webs on Earth. Using a minimal set of macromolecular scalings for a cell size distribution imputed from genomic data, we nevertheless recapitulate a biologically realistic ecosystem elemental composition.

Our imputed particulate N:P ratio is on the order of magnitude of the most generous ammonium to phosphate ratio Postberg et al. [55] suggest. Their study estimates N:P ratios from 10^6^ to 10 based off of a reported possible ammonium concentration of approximately 0.1 M and potential phosphate concentrations ranging from 10^*−*7^ *−* 10^*−*2^ M [55, 68, 72]. These numbers suggest phosphate concentrations ranging from comparable to much higher than predicted by a geochemical modeling study [23]. Both of these studies posit that phosphate would be at a high enough concentration to not be limiting for life, given that their reported concentrations are orders of magnitude higher than those observed in Earth’s surface ocean [44]. However, due to the ratio of inorganic nitrogen to phosphorus ostensibly available for uptake (the supply ratio), we argue that our theoretical methanogen community would draw down phosphorus and become phosphorus-limited with excess nitrogen. Only at the highest suggested phosphorus concentrations would the community exhibit signatures of co-limitation or limitation by a secondary ‘trace’ nutrient (for example, iron in Earth’s ocean [44, 68]). Due to the high concentrations of inorganic nutrients predicted and the results of our chemostat study, energy limitation may also explain high residual concentrations of phosphate despite its stoichiometric imbalance with nitrogen. However, this assessment is conditioned on the assumption that putative Enceladus cells would have macromolecular components with elemental compositions that directly mirror terrestrial life. We can extend our analysis by considering alternative macromolecular and elemental building blocks and conducting a more agnostic inquiry about the viability of organisms on Enceladus.

### Relaxing the Assumptions of Terrestrial Biochemistry

Instead of calculating emergent N:P allometric relationships from empirically-fitted macromolecular models, we can increase generality by simulating ecosystems across a broad range of putative N:P scaling exponents. Along these lines, we also can relax the assumption of Redfield 16:1 N:P ratios for ‘average’ organisms of median size [24, 28] and scan across a range of potential equivalent characteristic stoichiometries. Finally, in the previous section, we imputed a cell-size distribution based on the genome lengths of known methanogenic Archaea. However, ecosystem-size distributions vary widely across Earth environments [9, 10], centered around a generality of power-law scaling of cell number proportional to the inverse of the cell volume, i.e., *N* = *v*^*−*1^. While this exponent has been associated with multiple fundamental biological mechanisms [7, 77], we can also relax this assumption and consider size distributions following other exponents.

Using a modeling approach described in Methods, we calculated community N:P ratio as a function of three variable parameters - an N:P allometric scaling exponent (Figure 3A y-axis), the intercept of the N:P scaling relationship (related to the characteristic ‘Redfield-like’ stoichiometry, Figure 3B y-axis), and the exponential slope of the cell size distribution (Figure 3 x-axes). Scanning over a grid range of all three parameters encompassing at least 2 orders of magnitude, we generated 497,699 different simulations of possible ecosystem N:P ratios (see Extended Data Table 3). These simulations cover N:P scaling exponents from -1 to 1, median N:P ratios from 1 to 100, and cell size-abundance scaling exponents from -3 (Earth-like size distribution) to 0 (equal numbers of cells independent of cell-size). Across all simulations, we found an average N:P ratio of 378 (sd=852), suggesting that Earth-like ecosystem ratios are among the more P-rich of our simulations. This result is influenced by the heavy tail of our modeled distribution, with a maximum observed N:P ratio of 7914. Indeed, approximately 34% of all modeled N:P ratios are less than or equal to 16, and the skewness and kurtosis of the simulated N:P distribution strongly deviate from a symmetrical (normal) distribution (skewness=3.76, kurtosis=19.8, Jarque-Bera normality test *p <* 1^*−*10^). These statistics suggest that varying the three scaling relationships of the exponent for cellular N:P as a function of volume, the intercept of the N:P-volume relationship, and cell size-abundance exponent, can produce large deviations from Earth-like stoichiometries even when centered around Earth-like parameter values.

**Figure 3.**
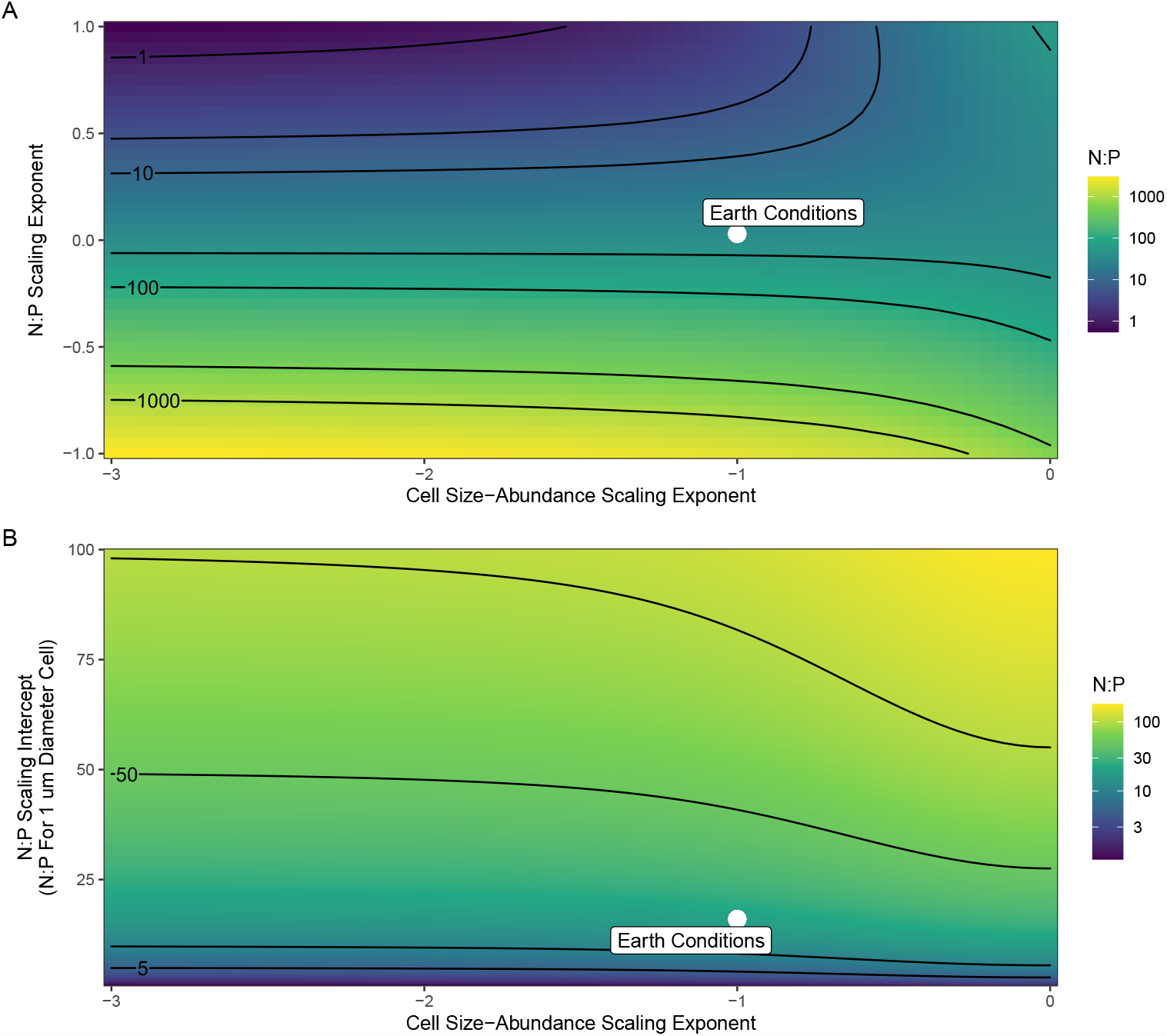
Heatmap of simulated ecosystem stoichiometries as a function of three free simulation parameters. The color in each cell represents the geometric mean of all simulations that use the combination of parameters specified by the associated x and y value. White points represent the simulated value of Earth-like parameters. A) Contours of ecosystem N:P as a bivariate function of cell size distribution scaling exponent and N:P scaling exponent. Contours represent N:P of 1, 5, 10, 50, 100, 500, and 1000. B) Contours of ecosystem N:P as a bivariate function of cell size distribution scaling exponent and intercept for the organismal N:P size-scaling relationship, using a 1 um cell as the reference cell size (see Methods). Contours represent 5, 10, 50, and 100.

This analysis suggests that using Earth-like scaling principles with non-Earth-like parameters can yield a wide range of possible ecosystem stoichiometries, some of which overlap with the dissolved N:P stoichiometries in Postberg et al. [55]. We analyzed which parameters had the greatest effect on ecosystem N:P (Figure 3). Comparing geometric-mean ecosystem N:P values across simulations varying two of the three free parameters at a time, we find that the N:P allometry has a large effect on ecosystem N:P compared to the cell size-abundance scaling exponent (Figure 3A), which only begins to strongly influence ecosystem N:P for slopes shallower than those observed on Earth (Figure 3A,B). We also find that holding all three parameters to Earth-like values does indeed recapitulate a ‘Redfield’ valued ecosystem stoichiometry (mean=15, sd=3.1, n=21 simulations), indicated by white point on 3A,B). In our simulations, 37.6% of ecosystems had N:P values within the range of [10^*−*6^, 10^*−*2^] posed for Enceladus inorganic N:P. This suggests that, if terrestrial assumptions about ecosystem size-structure and allometry are relaxed, the same scaling phenomena can produce ecosystems with Enceladus-like N:P ratios with a non-negligible probability, even using parameters centered around terrestrial values. Given that our distribution of simulated N:P varies over nearly 6 orders of magnitude, we do note that it is likely that terrestrial macromolecules with terrestrial average elemental compositions may not be reflective of the communities. Only 3.05% of simulations with an ecosystem N:P in the Enceledean inorganic N:P range use an N:P allometric exponent in the range suggested by the analysis of methanogen elemental scaling, while 69.1% use smaller exponents. This result suggests that cells comprising ecosystems with stoichiometries comparable to putative Enceledean stoichiometries are likely to use P in macromolecules that scale much more slowly than DNA, RNA, and phospholipids do in terrestrial organisms. Therefore, the N:P ratio of a putative Enceledean ecology may not correspond to N:P for Earth ecosystems. We then asked which pairs of elements constituent in terrestrial life have more similar ratios to those common in our simulations, and how those elemental pairs compare between cellular composition and environmental abundance.

### Applying Generalities in Terrestrial Biogeochemistry to Assess the Potential Biochemical Relevance of Phosphorus on Enceladus

The Earth’s ocean is a vast and highly heterogeneous ecosystem numerically dominated by microbial life [44]. Therefore, we used canonical ocean-wide average values of dissolved elemental concentrations and phytoplankton cellular elemental requirements for growth from [44] to analyze the relationship between planetary-scale biological and dissolved inorganic stoichiometries. Using the ratios of all non-redundant combinations of 17 different elements (see Methods for list of elements, Extended Data Table 4 for details) found in phytoplankton cells and seawater, we fit a scaling relationship between all pairs of cell quotas and their corresponding ratios in seawater (Figure 4A). We determined a significant sublinear scaling (cell quota= 41.7 *∗* 10^0.272*∗*seawater ratio^, 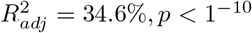). This suggests that there is an empirical basis on Earth’s oceans to relate dissolved stoichiometries to cellular stoichiometries.

**Figure 4.**
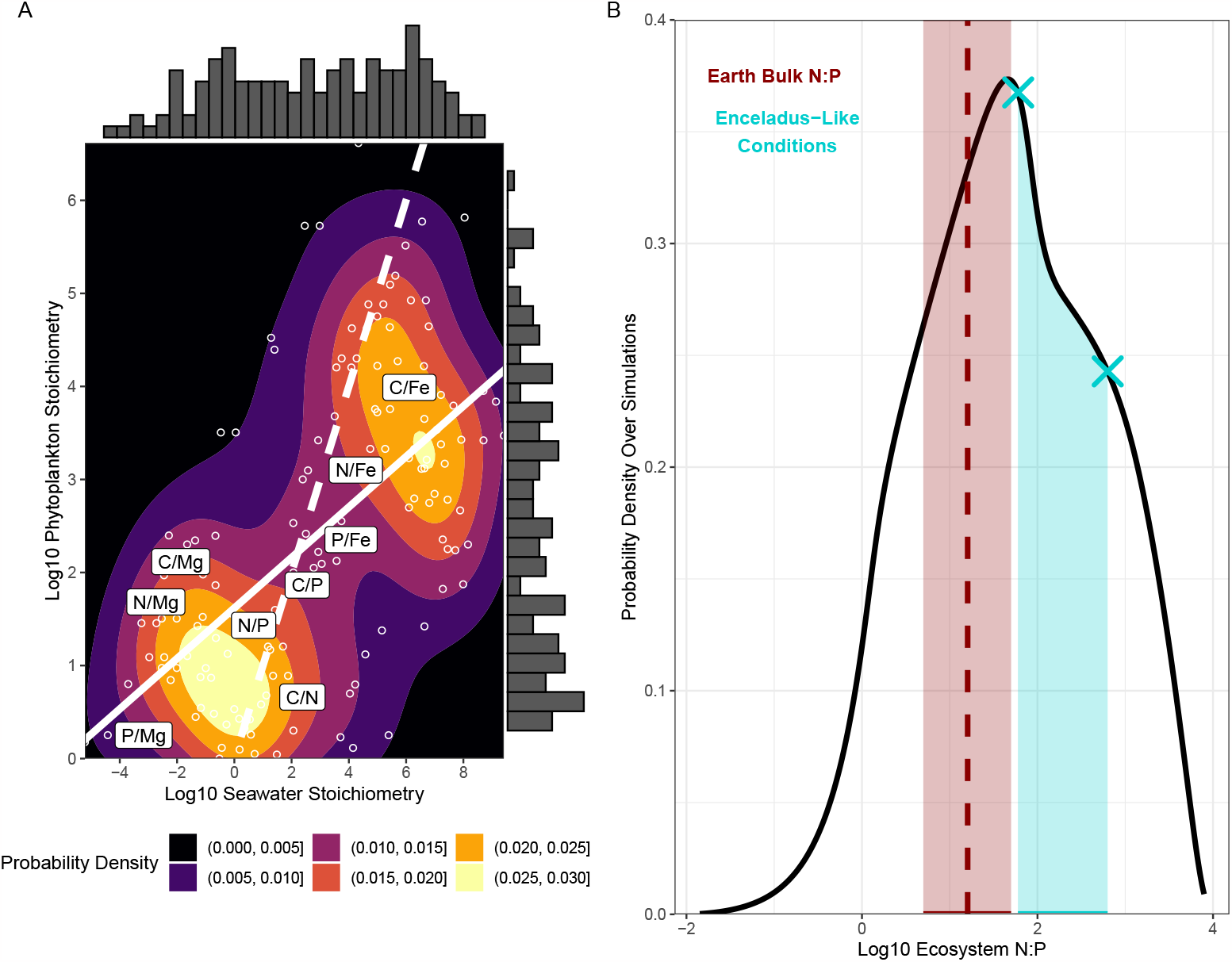
Scaling analysis of dissolved against biological elemental ratios and compatibility of predicted Enceladus stoichiometries with free parameter search. A) Observed correlation between ratios of elements dissolved in seawater (x-axis) and those same ratios in phytoplankton biomass (y-axis) across Earth’s oceans. White dashed line indicates 1:1 (note log scales), and solid white line indicates scaling line of best fit (exponent=0.272). Color denotes probability density in the bivariate distribution of elemental ratios, with marginal histograms denoting density of ratios for each axis, respectively. Example elemental ratios are labeled. B) Results from scaling analysis with relaxed assumptions about terrestrial parameters. Solid line indicates the probability density function (PDF) of ecosystem N:P ratios (note log scale) over the ensemble of approximately 500,000 simulated ecosystems varying relevant scaling parameters. The red dashed line indicates the Redfield ratio with the red shading representing observed variability for marine particulate organic matter from Station ALOHA in the North Pacific [27] The blue shading represents predicted ecosystem N:P ratios using the scaling relationship determined in (A) for proposed Enceledean stoichiometries. Blue crosses denote the likelihood values on the PDF corresponding to the minimum and maximum phosphate concentrations predicted in Postberg et al [55].

**Figure 5.**
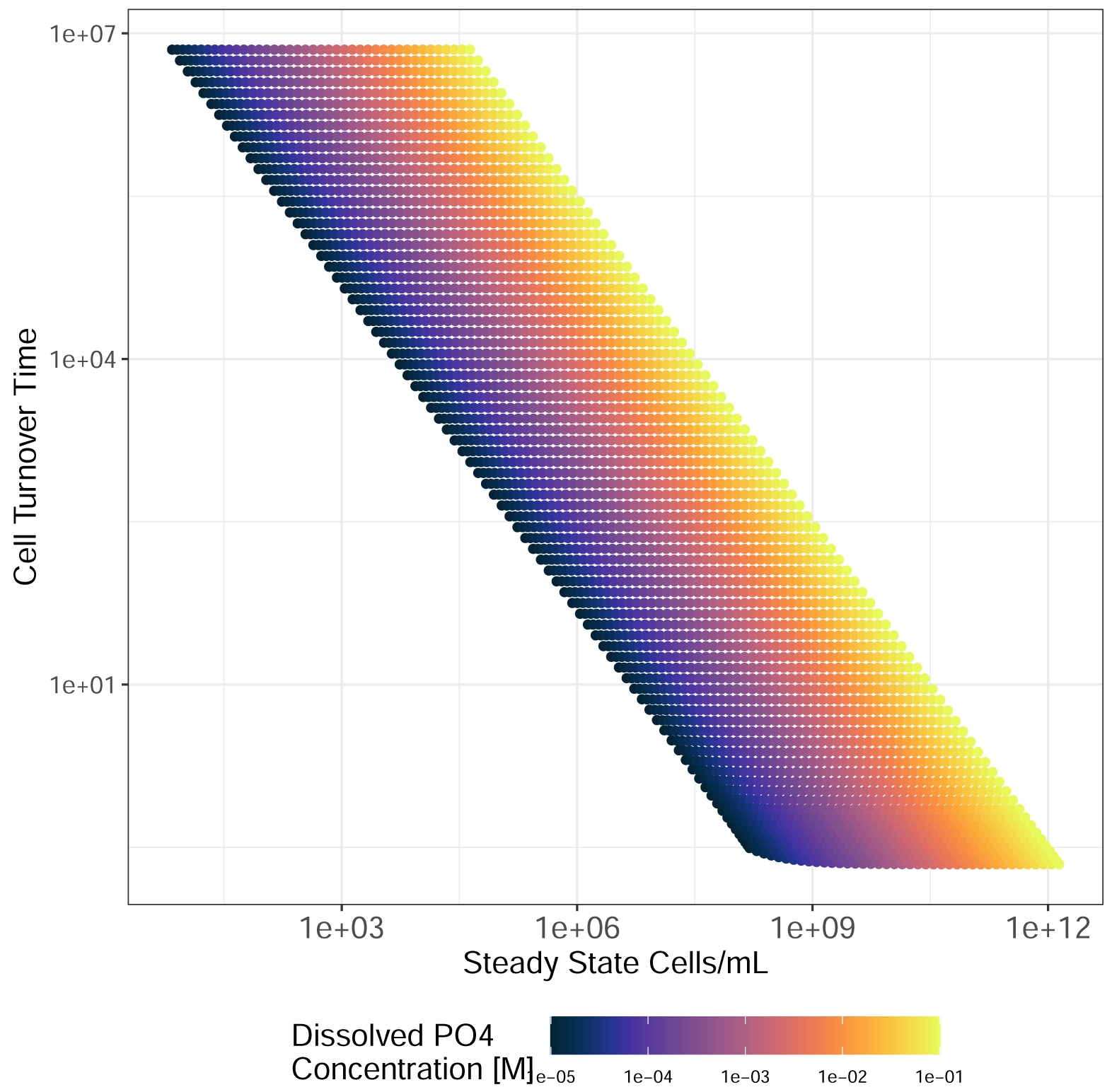
Extended Data Figure 1. Analysis of derived biomass turnover times to facilitate population stability for given steady state nutrient concentrations. For all chemostat model simulations, steady state dissolved phosphate concentrations are set by calculating the cell turnover time (chemostat washout rate) that corresponds to that phosphate concentration. Cell turnover time is shown on the y-axis while steady state biomass in cells/mL is shown on the x-axis. Point color indicates the steady state phosphate concentration set for that model simulation.

Using this model, we calculated what ecosystem N:P to expect from the range of dissolved N:P possible for Enceladus. We find that the 4 orders of magnitude in dissolved N:P collapse into a predicted range of biomass N:P ranging from 10^1.9^ to 10^3^, or approximately 79 to 1000 (see Extended Data Table 5). These ratios again far exceed average N:P values for Earth aquatic and marine ecosystems, supporting the conclusion that an Enceladus cellular N:P allocation would likely fundamentally differ from a terrestrial organism. We synthesized this data-driven analysis of data from Earth’s oceans with our parameter search simulations to calculate the probability of an arbitrary ecosystem having an N:P ratio compatible with the dissolved N:P ratios suggested for Enceladus. We find a probability of 0.297, suggesting a non-negligible probability of occurrence despite a drastic departure from Earth-like N:P ratios (Figure 4B). We note the parallel that through completely independent modeling perspectives, our calculated probability of an Enceladus dissolved N:P ratio being consistent with an arbitrary ecosystem matches the probability of a living methanogenic ecosystem of 0.32 reported in Affholder et al. [1].

We then compared the ratios suggested by our scaling model to the observed cell quotas from [44] to identify which pairs of elements in phytoplankton cells have similar relationships to a hypothetical Enceladus cell’s N:P. We found that these ratios most closely resemble C:S or C:P (95 and 124, respectively) at the lower end and combinations of S, P, and major cations (Ca, K, Mg) against trace transition metals (Mn, Ni, Zn) at the higher end. This result suggests that if corresponding scaling relationships between environmental and biological ratios are applicable to Enceladus, hypothetical Enceladus cells would use phosphorus as a trace constituent.

## Discussion

### Stoichiometry as a Tool to Probe Exotic Environments for Ecological Compatibility

Here we use multiple modeling approaches all centered around the meta-framework of considering the relative ratios of different elements in an environment as a biosignature. Our chemostat model, which leverages a few free parameters (energetic limits on maximum growth rate, washout rate - see Methods) to calculate steady state cell density as a function of steady state phosphorus concentration, predicted ranges of supportable cell densities comparable to those previously offered for the Enceladus ocean [52, 64, 69] at high phosphate concentrations, but at the low end of the range of phosphate concentrations suggested by Postberg et al. [55], we predict lower supportable cell densities - potentially a signature of phosphate limitation at these low concentrations. We used genomic data on methanogenic Archaea to predict the macromolecular composition of an ecosystem comprised of these organisms, and found that our estimated ecosystem N:P ratio matches, within an order of magnitude, the ammonium:phosphate ratio implied by the maximum concentration from Postberg et al. [55]. Expanding to a more agnostic view of N and P incorporation into biomass, we found that integrating elemental allometric scaling laws for cells and ecosystem cell size distribution scaling laws could recapitulate the full range of possible N:P ratios implied by Postberg et al. [55], taking up approximately 30% of the parameter space we simulated - matching estimated likelihoods of Enceladus as habitable for methanogens from a Bayesian perspective based on plume composition reported in Affholder et al. [1]. Taking these results together, we conclude that recent reports of phosphorus in the Enceladus plume suggest conditions marginally compatible with the potential to support an ecosystem. Notably, our analysis is not an attempt at life detection, nor do we argue that the results of our analysis are strong evidence as to whether or not the reported phosphorus values constitute a biosignature. Rather, we aim to illustrate the utility of matter-oriented ecological and physiological modeling to complement energy-oriented metabolic models as a method to interpret compositional data about extraterrestrial environments with respect to life detection. In addition, we think these results do not rule out the possibility of life on Enceladus.

The framework and suite of methods we present here are useful in part because of their wide generality. The same modeling procedures we use for this analysis can be readily adapted to consider elemental or molecular ratios in any kind of exotic environment - from Enceladus to Europa, Mars, or even deep time in Earth history prior to robust molecular fossil records. This stands in contrast to metabolism-based models, which would need to be completely reframed to consider relevant redox pairs and thermodynamic conditions in each new environment.

### Possible Limits to Life on Enceladus

Previous geochemical modeling studies analyzing the potential rates of methanogenesis and metabolism of putative Enceladus life have suggested imperfect total conversion of hydrogen and carbon dioxide into methane, leaving a residual energy signature [26], while other modeling efforts have suggested that total cell counts in hydrothermal vent communities would higher than average population densities on Earth if all available substrates for metabolism were consumed [69]. However, it is possible that even in the low-energy circumstances of dark, anaerobic methanogenesis as the mode of primary production (compared to oxygenic photosynthesis on Earth [12]), resources to synthesize biomass may be limiting to life [40, 60]. While the total estimated standing stocks of inorganic nitrogen and phosphorus appear higher than in Earth’s oligotrophic oceans on Enceladus, our study suggests that phosphorus may be limiting to Earth-like cells under most of the predicted parameter regime. High standing stocks of these nutrients could be consistent with incomplete draw down due to a small or metabolically slow biosphere, a biosphere with a recent origin of life, inefficient substrate uptake affinities, or high cell turnover or mortality rates resulting in an imbalance between remineralization and biological fixation of these elements.

Beyond terrestrial macronutrients like nitrogen and phosphorus, it is also possible for putative life on Enceladus to be limited by bioactive trace elements. In a laboratory culture study, Taubner et al. suggest the nickel cofactor in methyl coenzyme M reductase, a necessary enzyme for Earth methanogenesis, may limit methanogenesis and suggest organic nickel complexes as a possible biosignature candidate [68]. This theory has also been extended to the relative rarity of molybdenum, another trace enzyme cofactor, in the solar system [40]. The same phenomenon of trace metal limitation for enzyme cofactors is detected in Earth’s oceans, where High Nutrient Low Chlorophyll (HNLC) regions occur due to an excess of macronutrients relative to trace metals [49]. However, we caution that these cofactors do not necessarily make for ‘smoking-gun’ biosignature candidates, as even terrestrial life has evolved a multiplicity of enzymes performing a common function using different sets of metal cofactors - superoxide dismutase among marine bacteria has evolved 4 different metal forms [76]. Better understanding the generality of trace metal enzyme cofactors in biochemistry and their geochemical dynamics will help to better incorporate them into a stoichiometric model-based life detection instrument.

The high concentrations of macronutrient elements, and particularly of ammonium, in Enceladus plume data have also raised the concern that life on Enceladus would have to overcome ammonium toxicity [42]. Ammonium resupply rates in excess of biological uptake may also prevent draw down and induce observed stoichiometric imbalances compared to Earth life. However, laboratory culture experiments using terrestrial methanogens subjected to Enceladus-like ammonium concentrations did not exhibit strong growth inhibition unless in the presence of other inhibitors, suggesting that ammonium toxicity may not be a relevant mechanism preventing the proliferation of life [68].

Through chemostat modeling, we find a range of possible supportable biomass estimates using Earth physiological parameters that straddle energy limitation and resource limitation. Our results fall within the ranges previously reported from metabolic modeling based off of methanogenic Archaea. However, we find that the maximum attainable cell densities at the minimum end of the range of reported phosphorus concentrations presented in [55] are smaller than estimates based on efficient consumption of the substrates required for methanogenesis - potentially suggesting phosphorus limitation. At the high end of the reported phosphorus concentrations, energy availability becomes much more important as our range of steady state cell densities span the full gamut of previously reported estimates [52, 64, 69]. As phosphate concentrations on Enceladus get increasingly precise estimates, this model could allow us to discriminate between a primarily energy-limited and nutrient-limited regime. There may be a fundamental energetic limit on the ability of life to do uphill chemical reactions, but that doesn’t preclude matter-based approaches in being informative particularly with respect to mass spectral data which are the most readily available for life detection missions.

### Reimagining Phosphorus as a Trace Constituent of Enceledean Cells

A previous study assessing the viability of terrestrial contamination of Enceladus found that adding phosphorus to the ‘seed set’ of bioactive molecules improved metabolic network expansion but was still insufficient to recapitulate an entire Earth cellular metabolism as inferred by genomic content [60]. Our models find that the predicted dissolved N:P ratio is more consistent with an ecology that scales cellular phosphorus content more slowly than terrestrial cells with volume, and therefore is more likely to use phosphorus in a smaller set of macromolecules than terrestrial life.

The possibility remains for life to use a completely novel set of macromolecules, completely forgoing membrane lipids or nucleic acids connected by phosphate bridges. However, this possibility is not necessary to explain life with drastically reduced phosphorus content, as macromolecular evolution on Earth has rendered multiple examples of standard macromolecules with the phosphorus replaced by other elements throughout its history. Recent research on early Earth life has suggested a ‘thioester world’ pre-phosphorus prebiotic chemistry, in which energy that is now stored in ATP by cells may have been stored in thioester molecules such as CoA [21, 48]. A network expansion study based on a small suite of likely prebiotic molecules on Earth excluding phosphorus was able to recapitulate many reactions required for energy metabolism, largely relying on thioester molecule-mediated reactions [21]. Similarly, biomedical technology has used disulfide-bridges to replace phosphate bridges in artificial nucleotide polymers [4, 13]. In phosphorus-limited regions of Earth’s surface ocean, there is a distinct macromolecular signature of phosphorus replacement with nitrogen-bearing lipids [70], suggesting that natural selection on Earth has identified mechanisms of phosphorus replacement in lipids. Such evolutionary and physiological adaptations manifest in the broad variance of biological N:P ratios among different microbial ecosystems around Earth [24, 59, 66, 67]. Therefore, it is not unreasonable to expect that alternative origins of life may have evolved to not use phosphorus as a constituent of many key macromolecules such as has transpired on Earth.

### Priorities for Continued Astrobiological Research on the Viability of Enceladus

Based on new analyses from the Cassini mission, we have conducted a preliminary, size-scaling informed, ecological stoichiometry modeling study. We assessed the likelihood of reported phosphorus concentrations with: 1) living ecosystems resembling terrestrial methanogens, 2) according to more general elemental ratio biogeochemical scaling, and 3) generalizing away from terrestrial macromolecular and/or elemental compositions. Our findings suggest that, while we did find marginal evidence for ecosystem stoichiometries comparable to reported Enceladus N:P stoichiometry, due to the high N:P ratios reported for the Enceladus ocean, hypothetical ecosystems would generally experience an N-heavy resource supply ratio, or cellular elemental composition would have a much lower P-content than life on Earth is generally known to. We suggest two priorities for further astrobiological research to better understand the implications of these conclusions.

First, we echo previous calls in the astrobiology literature to explore more generalized notions of metabolism and physiology [28, 42, 51]. Our analyses suggest that while assuming terrestrial allometry and ecosystem size-structure could yield theoretical non-negligible biomass and biological stoichiometries that are within the ranges of N:P ratios suggested by Cassini data, these cases are marginal. More of the parameter space of possible ecosystems consistent with the Cassini data implies macromolecular and elemental architecture that differs from those common on Earth. We suggest that looking for direct parallels to terrestrial life in the form of biochemistry may then be ill-informed as a life detection strategy for Enceladus. However, to maintain principled life detection hypotheses, a more detailed understanding of alternative, yet viable, cellular architectures and ecosystem trophic structures are possible - particularly given our results that phosphorus may only be a trace constituent of biomass and that ecosystem size-structure may substantially deviate from standard power-law scaling on Earth.

Second, we recommend broadening the scope of Earth analogue environments to include those with extreme resource supply ratios mirroring that suggested for Enceladus. Previous studies have offered selection criteria for Earth analogues of ocean worlds and other life detection candidates have centered around energetic parallels, such as light availability, temperature, carbon sources, and sources of chemical energy [14, 20, 42]. In particular, Earth environments such as deep subsurface sediments, the Lost City hydrothermal vent system, and cold gas hydrates [20, 42, 69]. Studies attempting to mirror Enceladus conditions to test the viability of terrestrial methanogens have also emphasized temperature, pressure, and relative supply of redox components as key variables [64, 68, 69]. Our study finds that in addition to these extreme energetic conditions, potential life on Enceladus would also face extreme resource supply ratios relative to most environments on Earth. Our scaling analysis suggests that an Earth-like N:P allometry is not very compatible with reported inorganic N:P ratios (using this scaling exponent only resulted in 3% of the Enceladus-like putative ecosystem ratios from our parameter search simulations). Therefore, to better understand potentially relevant biomolecules, it would behoove us to further characterize the variety of P-reduced biomolecules that have been discovered by natural selection on Earth. One advantage is that such environments are potentially more accessible than famous analog environments such as the Lost City vent system - Lake Superior of the Laurentian Great Lakes is one of the few aqueous environments on Earth with dissolved inorganic N:P ratios approaching the possible 10^6^ reported in Postberg et al. [55], being as high as 10^4^ [16, 65]. The Sargasso sea is another surface ocean environment with persistently high dissolved N:P ratios, and studying microorganisms adapted to that P-limited environment has broadened our understanding of lipid dynamics and microbial carbon cycling [41, 45, 62, 63, 70]. Having a better understanding of the macromolecular evolutionary dynamics in these persistently P-limited microbial ecosystems may then improve the scope of possible alternative physiologies that may match low-biological-P ecological hypotheses such as the one presented in this study.

## Methods

### Chemostat Modeling

Chemostat models have been used as the simplest representation of biogeochemical systems. These models consider the influx of inorganic nutrients and energy sources, uptake and transformation of nutrients into cellular materials as a function of cellular grow, and the loss of cells, nutrients, and energy sources from the system as function of efflux. Such models are typically written for each nutrient as

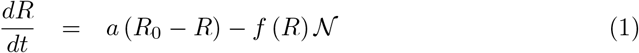

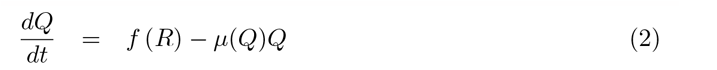

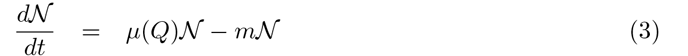

where *µ*(*Q*) is the growth rate as a function of all of the existing elemental quotas and energy sources (cellular quantities), and is typically given by

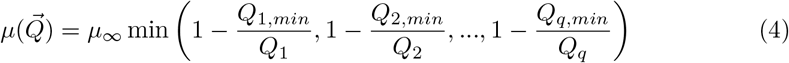

where the minimum is taken over the total number of limiting factors and one can write down Equations 1 for each nutrient [31–34, 37].

In this model *R*_0_ is the concentration outside the system, *a* is the flow rate of the system, *m* and the loss of the nutrients from the system including efflux and cellular mortality from a variety of factors. Typically we take *m* = *a* for the simplest scenarios. In this framework all of the parameters are determined by cell size [28], and *f* typically takes the form

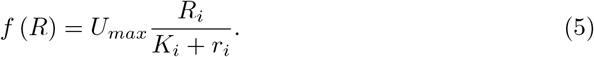

These equations can be solved for the steady-state values of the environment and cellular states giving

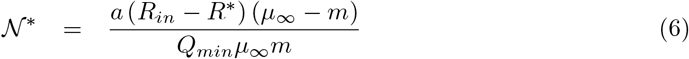

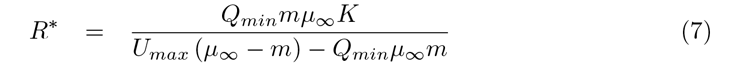

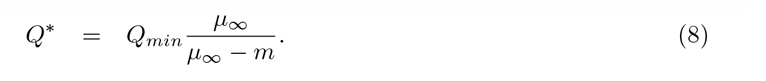

[31–34, 37] For our analysis we took a typical bacterial cell volume of 10^*−*18^ m^3^ to determine the cellular parameters using the allometric relationships in [28], and we considered *a* = 0.007 day^*−*1^. We scaled maximum growth rate by the energy availability. Then for each *R*^*∗*^ we used the steady state equations to solve for *m* and we used this *m* combined with *R*^*∗*^ to calculate the *N* ^*∗*^.

### Methanogen Genome Data Processing

Data on genome lengths of sequenced methanogens were recovered from supplementary data from [56]. Only complete genomes were retained for further analysis, as indicated by the designation ‘complete assembly’ in the data file. Initially, to reflect likely low-temperature conditions on Enceladus [55], we screened for methanogens labelled as ‘psychrotolerant’. Only 8 out of 74 available complete genomes were marked as psychrotolerant, so we compared the genome lengths of psychrotolerant taxa to the overall genome length distribution and found no deviation from the overall pattern (Supplementary Figure [X]).

### Cell Volume and Cell Size Distribution Estimation

All computations were conducted using R v4.2.1 [57]. To do the single conversion from genome length (in base pairs) to cell volume, we inverted the power law scaling presented in [29]. This reverse scaling was implemented for all observed genome lengths in the dataset to make an estimated cell volume distribution. Because there were only 74 complete genomes, we aimed to infer the overall size distribution to reduce potential bias from undersampling in the observations. From the distribution of inferred cell volumes, we fit a gamma distribution using the MASS library function fitdistr [71]. A gamma distribution was chosen due over an exponential or power-law distribution due to the truncation at small volumes and was chosen over a symmetric distribution due to the heavy tail. Noticing possible multimodality in the volume distribution, we also assessed whether or not fitting a mixture of gamma distributions improved the overall fit by using the mixtools library [8]. We found no substantial increase in log likelihood using a mixture of gamma distributions and visual inspection of the fitted model showed no improvement in fit. Log-spaced bins were sampled from the fitted gamma distribution to simulate the theoretical cell volume distribution.

### Macromolecular and Stoichiometric Modeling

From the inferred cell volume distribution, we applied macromolecular scaling laws converting cell volume to pg protein, pg lipids, pg carbohydrates as reported in [17] using the mean exponent and intercept for the power law fit to all aggregated taxa. We used the same genomic power law as in the above section and the power-law ribosomal scaling from [29], and converted *m*^3^ of each macromolecule type into pg using the average mass of a nucleotide (617.96 g/mol nucleotide) and the density of ribosomes (1.42 *∗* 10^6^ *g/m*^3^) from [citations] to convert to pg macromolecule.

We used the elemental compositions of each macromolecular type as reported in [19] to convert pg macromolecule to pg C,N, and P. For ribosomes, we used a ratio of 38% protein and 62% RNA as reported in [43] bionumber ID:109047. For lipids, we used an ensemble average of the reported ranges of the relative proportion of phospholipids to non-phospholipids reported in [19]. While phospholipid subsitution is a common adaptation to overcome low-phosphorus environments [70], the contribution of lipids to total cellular phosphorus content only becomes apparent at the largest of the cell volumes we analyze here (Supplemental Figure [X]), so we find our overall assessment to be robust to variation in this proportion.

Total cellular stoichiometry was calculated by summing the pg of C, N, and P for each macromolecular type for a give cell volume. We calculated ecosystem stoichiometry as a weighted average over all analyzed cell volumes of the total cellular pg C/N/P, using our fitted gamma cell volume distribution as the weights. A power-law was fit to the allometric N:P scaling using the nls function from the R base stats library.

### Ecosystem Stoichiometry Parameter Search

Based on the scaling principles described in the previous section, we developed a generalized scaling framework to estimate ecosystem N:P based on the generic model:

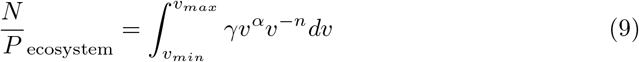

Where *v* is the cell volume, *α* is a scaling exponent that represents the size-scaling of the N:P ratio of a single cell based on its macromolecular components, *γ* is the intercept of the N:P size-scaling relationship (determined by the ‘Redfield’-like average cell stoichiometry), *n* is the exponent representing the steepness of the cell size distribution (the higher the value of *n* the rarer large cells are in the ecosystem compared to small cells), and *v*_*min*_ and *v*_*max*_ represent the minimum and maximum cell volume in the ecosystem. In words, the ecosystem N:P ratio is the sum of the N:P ratios for each individual cell size weighted by its relative abundance in the population.

The intercept parameter *γ* was determined by assuming a characteristic stoichiometry for a cell of median volume in the size distribution (in our simulations taken to be 10^*−*18^*m*^3^ or approximately the size of *E*.*coli*), then fitting a given N:P size-scaling exponent to a power-law relationship such that the value of cellular N:P at this standard volume matched the characteristic stoichiometry. For a given N:P scaling exponent, *α*^*∗*^, and a given characteristic stoichiometry, *S*^*∗*^, The model was solved as follows:

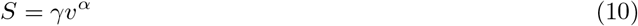

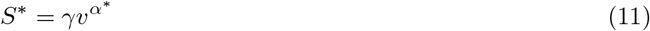

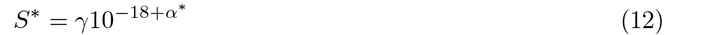

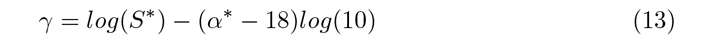

Given the ubiquity of size-abundance scaling exponents with volume close to -1 in phytoplankton cell size distributions [9], we varied the value of *n* from -3 (much more biased to small cells than Earth oceans) to 0 (no bias toward small or large cells). According to principles of trophic transfer and sublinear scaling of surface area to volume ratios for inorganic substrate uptake, we did not consider environments where the size distribution would scale with a positive exponent. Then, based off of the scaling relationship of N:P based on the macromolecular scaling laws used in the previous section having a very slight positive value (*≈* 0.03), we searched across values of *α* going from -1 to 1 in linear spaced bins of 0.05 to consider the Earth-like scaling relationship as ‘average’, while also considering cases where cellular P content scales much faster than cellular N content (negative values) and cases where cellular N content scales much faster than cellular P content (positive values). Finally, we considered median cellular stoichiometries that varied by 1 order of magnitude on either side fo the Redfield ratio for Earth (1-100) to recalculate *γ* for each *α* in linear-spaced bins of 0.5. Altogether, this led to 497,699 unique parameter combinations leading to different estimated ecosystem N:P ratios. Statistics were calculated using the R v4.2.1 [57] base functions and the moments package [35] to calculate skewness and kurtosis and to conduct statistical tests for heavy-tailedness.

### Earth Cell-Quota to Seawater Stoichiometric Scaling

Data from Moore et al. [44] Supplemental Table 1 were compiled and elements were ordered by decreasing relative proportion in phytoplankton biomass (cell quota relative to carbon) - with the exception of hydrogen and oxygen given their dominant concentrations in seawater. All 17 remaining elements with reported average dissolved concentrations in seawater and phytoplankton cell quotas (C, N, Si, K, S, P, Mg, Ca, Fe, Sr, Mn, Ni, Zn, Cu, Cd, Co, Mo) were then turned into 136 ratios of both cellular quota and seawater stoichiometry (with the cellular quota always greater than 1 and no redundant pairs 0 136 is equal to 17 choose 2) for further analysis.

A power-law relationship between seawater ratio and phytoplankton cell quota was then estimated using ordinary least squares on the log10 values of the data, resulting in a relationship of *log*10(*cell*) = 1.622 + 0.273*log*10(*seawater*) with 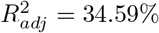 and *p <* 1^*−*10^. The same relationship was then estimated using a bayesian linear regression approach using the R v4.2.1 [57] package BGLR [50] for both a null model without the seawater ratio as a predictor and with the seawater ratio as a predictor. The model with the seawater predictor converged to similar parameter estimates as the ordinary least squares model and had higher adjusted likelihood compared to the null model, further suggesting the model as a good fit. We also conducted major axis regression on the same relationship using the R package lmodel2 version 1.7-3 [36] to check for robustness and found a similar significant relationship (permutation tests, p¡1e-4) with similar parameter values using this estimation procedure.

The scaling law was then used to predict the cell quotas for the range of seawater stoichiometries for N:P reported in [55]. We used these predictions as ranges to calculate the probability density of ecosystems having stoichiometries in the distribution of simulated N:P values as described in the above section, and arrived at a probability density of 0.297.

## Data and Code Availability

Data and code necessary to reproduce the analyses and figures for this paper are available at https://github.com/dmuratore/enceladus_stoichiometry with an archived DOI at https://zenodo.org/doi/10.5281/zenodo.10045287.

## Supporting information

Captions for Extended Data Tables and Figures

## Acknowledgments

C.P.K, S.I.W, H.G., and C.H.H were supported by Agnostic Biosignatures award no. 80NSSC18K1140. D.M. was supported by the Santa Fe Institute and the Omidyar Fellowship.

## Extended Data

Extended Data Table 1. Simulation results from chemostat model. Simulations searched for steady state biomass in cells/mL by varying the relative energy efficiency of phosphate conversion into biomass (first column), the steady state dissolved phosphate concentration (second column), and using those two parameters, calculating the necessary chemostat outflow/cell turnover rate to allow ecosystem stability (fourth column). The steady state biomass for each simulation is in the third column.

Extended Data Table 2. Power law parameters for macromolecular allometric scaling models. Macromolecules calculated include total genome, total proteins, total ribosomes, total lipids, and total carbohydrates to represent the largest pools of intracellular C, N, and P. Parameters are presented in the form of a log-log relationship, i.e., *log*_10_(picograms macromolecule) = *b* + *alog*_10_(cell volume). The slope parameter, *a*, is in the power-law slope column, while the imntercept parameter, *b* is in the power-law intercept column. References for each scaling are provided.

Extended Data Table 3. Results from scaling parameter grid search analysis. Each row represents one of 497,699 simulated ecosystem N:P ratios (first column) based on the model described in Methods. The parameters used for these calculations are the cellular allometric scaling exponent for N:P ratio as a function of cell volume (second column), the intercept for the N:P-volume relationship (third column), which is determined by setting a “Redfield-like” average N:P stoichiometry for a cell with a 1micron radius (fourth column). That scaling is integrated over a power-law cell size distribution governed by a scaling exponent (fifth column), where the cell volume distribution on Earth is characterized on average by an exponent of -1 and an exponent of 0 refers to an ecosystem with equal abundances of all cell volumes.

Extended Data Table 4. Data compiled from Moore et al. (see Methods) on average phytoplankton cell quotas and average dissolved seawater concentrations of 17 different bioactive elements (see Methods for enumerated list). Elements are presented as ratios, with the more abundant element in phytoplankton biomass always as the numerator. The columns show the ratio (first column), the numerator element (second column), the denominator element (third column), the average phytoplankton biomass stoichiometry (fourth column), and the average dissolved seawater stoichiometry (fifth column). Rows are ordered by the most abundant element in phytoplankton for the ratio numerator, then denominator.

Extended Data Table 5. Scaling-based estimates of ratios from major elemental pairs highlighted in main text figure 4. Columns show the elemental ratio (first column), the dissolved seawater ratio (second column), the scaling-model based estimate of the phytoplankton biomass stoichiometry with 95% confidence intervals in parentheses (third column), and then the observed phytoplankton biomass ratio from Extended Data Table 4 (fourth column). For Enceladus-based estimated, the observed biomass ratios are listed as NA.

